# Multi-Connection Pattern Analysis: Decoding the Representational Content of Neural Communication

**DOI:** 10.1101/046441

**Authors:** Yuanning Li, R. Mark Richardson, Avniel Singh Ghuman

## Abstract

The lack of multivariate methods for decoding the representational content of interregional neural communication has left it difficult to know what information is represented in distributed brain circuit interactions. Here we present Multi-Connection Pattern Analysis (MCPA), which works by learning mappings between the activity patterns of the populations as a factor of the information being processed. These maps are used to predict the activity from one neural population based on the activity from the other population. Successful MCPA-based decoding indicates the involvement of distributed computational processing and provides a framework for probing the representational structure of the interaction. Simulations demonstrate the efficacy of MCPA in realistic circumstances. Applying MCPA to fMRI data shows that interactions between visual cortex regions are sensitive to information that distinguishes individual natural images, suggesting that image individuation occurs through interactive computation across the visual processing network. MCPA-based representational similarity analyses (RSA) results support models of error coding in interactions among regions of the network. Further RSA analyses relate the non-linear information transformation operations between layers of a computational model (HMAX) of visual processing to the information transformation between regions of the visual processing network. Additionally, applying MCPA to human intracranial electrophysiological data demonstrates that the interaction between occipital face area and fusiform face area contains information about individual faces. Thus, MCPA can be used to assess the information represented in the coupled activity of interacting neural circuits and probe the underlying principles of information transformation between regions.

## Introduction

Since at least the seminal studies of Hubel and Wiesel (Hubel and Wiesel, 1959) the computational role that neurons and neural populations play in processing has defined, and has been defined by, how they are tuned to represent information. The classical approach to address this question has been to determine how the activity recorded from different neurons or neural populations varies in response to parametric changes of the information being processed. Single unit studies have revealed tuning curves for neurons from different areas in the visual system responsive to features ranging from the orientation of a line, shapes, and even high level properties such as properties of the face (Desimone et al., 1984; Hubel and Wiesel, 1959; Tsao et al., 2006). Multivariate methods, especially pattern classification methods from modern statistics and machine learning, such as multivariate pattern analysis (MVPA), have gained popularity in recent years and have been used to study neural population tuning and the information represented via population coding in neuroimaging and multiunit activity (Cox and Savoy, 2003; Ghuman et al., 2014; Haxby et al., 2001; Haynes and Rees, 2006; Hirshorn et al., 2016; Kamitani and Tong, 2005; Poldrack, 2011; Polyn et al., 2005). These methods allow one to go beyond examining involvement in a particular neural process by probing the nature of the representational space contained in the pattern of population activity (Edelman et al., 1998; Haxby et al., 2014; Kriegeskorte and Kievit, 2013).

Neural populations do not act in isolation, rather the brain is highly interconnected and cognitive processes occur through the interaction of multiple populations. Indeed, many models of neural processing suggest that information is not represented solely in the activity of local neural populations, but rather at the level of recurrent interactions between regions (Grossberg, 1980; Kveraga et al., 2007; Lee and Mumford, 2003). However previous studies only focused on the information representation within a specific population (Freiwald et al., 2009; Ghuman et al., 2014; Haxby et al., 2001; Hirshorn et al., 2016; Nestor et al., 2011; Tsao et al., 2006), as no current multivariate methods allow one to directly assess what information is represented in the pattern of functional connections between distinct and interacting neural populations with practical amounts of data. Such a method would allow one to assess the content and organization of the information represented in the neural interaction. Thus, it remains unknown whether functional connections passively transfer information between encapsulated modules (Fodor, 1983) or whether these interactions play an adaptive computational role in processing. Note that in this context non-adaptive information transfer is equivalent to a static linear projection where no computational “work” is done in the interaction between the regions and therefore no information is added. Adaptive information transfer is one in which computational work related to the behavioral state or condition is performed and therefore state or condition specific information is added through the interaction between regions; this is equivalent to a non-linear function.

Univariate methods that go beyond assessing the degree of coupling between populations to assess changes in the relationship between the activity as a factor of condition also examine adaptive communication between regions. For example the psychophysiological interactions (PPI; (Friston et al., 1997)) and dynamic causal modeling methods (Friston et al., 2003) are sensitive to adaptive interregional communication. However, when compared with univariate methods, it has been noted that multivariate methods allow for “more sensitive detection of cognitive states,” “relating brain activity to behavior on a trial-by-trial basis,” and “characterizing the structure of the neural code” (Norman et al., 2006). Thus, a multivariate pattern analysis method for functional connectivity analysis is critical for decoding the representational structure of interregional interactions.

In this paper, we introduce a multivariate analysis algorithm combining functional connectivity and pattern recognition analyses that we term Multi-Connection Pattern Analysis (MCPA). MCPA works by learning the discriminant information represented in the shared activity between distinct neural populations by combining multivariate correlational methods with pattern classification techniques from machine learning in a novel way. Much the way that MVPA goes beyond a t-test or ANOVA by building a multivariate model of local activity that is then used for single-trial prediction and classification, MCPA goes beyond PPI by building a multivariate connectivity model that is then used for single-trial prediction and classification. This single-trial prediction and classification makes MCPA distinct from previous connectivity approaches that only statistically test the absolute or relative functional connectivity between two populations (Cribben et al., 2012; Finn et al., 2015; Richiardi et al., 2011; Shirer et al., 2012; Wang et al., 2015) and allows for a detailed probe of the representational structure of the interaction.

The MCPA method consists of an integrated process of learning connectivity maps based on the pattern of coupled activity between two populations A and B conditioned on the stimulus information and using these maps to classify the information representation in shared activity between A and B in test data. The rationale for MCPA is that if the activity in one area can be predicted based on the activity in the other area and the mapping that allows for this prediction is sensitive to the information being processed, then this suggests that the areas are communicating with one another and the communication pattern is sensitive to the information being processed. Thus, MCPA simultaneously asks two questions: 1) Are the multivariate patterns of activity from two neural populations correlated? (i.e. is there functional connectivity?) and 2) Does the connectivity pattern change based on the information being processed? This is operationalized by learning a connectivity map that maximizes the multivariate correlation between the activities of the two populations in each condition. This map can be thought of like the regression weights that transform the activity pattern in area A to the activity pattern in area B (properly termed “canonical coefficients” because a canonical correlation analysis [CCA] is used to learn the map). These maps are then used to generate the predictions as part of the classification algorithm. Specifically, a prediction of the activity pattern in one region is generated for each condition based on the activity pattern in the other region projected through each mapping. Single trial classification is achieved by comparing these predicted activity patterns with the true activity pattern (see Figure 1 for illustration). With MCPA single trial classification based on multivariate functional connectivity patterns is achieved allowing the nature of the representational space of the interaction to be probed.

**Figure 1.**
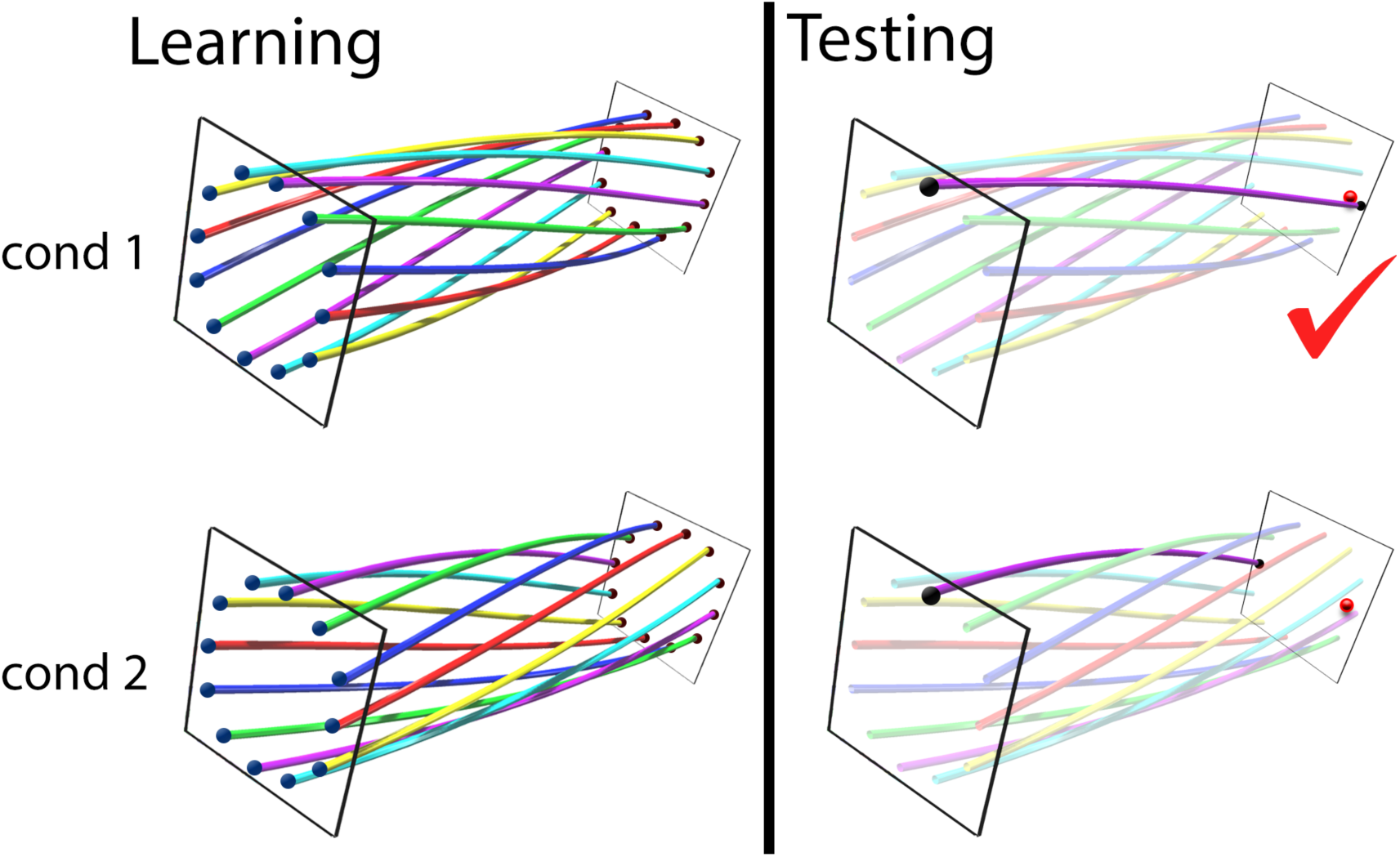
Illustration of the connectivity map and classifier of MCPA. The MCPA framework is demonstrated as a two-phase process: learning and testing. **Top left:** An illustration of the learned functional information mapping between two populations under condition 1. The representational state spaces of the two populations are shown as two planes and each pair of blue and red dots correspond to an observed data point from the populations. The functional information mapping is demonstrated as the colored pipes that project points from one space onto another (in this case, a 90 degree clockwise rotation). **Bottom left:** An illustration of the learned functional information mapping between two populations under condition 2 (in this case, a 90 degree counterclockwise rotation). **Top right:** An illustration of the predicted signal by mapping the observed neural activity from one population onto another using the mapping patterns learned from condition 1. The real signal in the second population is shown by the red dot. **Bottom right:** An illustration of the predicted signal by mapping the observed neural activity from one population onto another using the mapping patterns learned from condition 2. In this case, MCPA would classify the activity as arising from condition 1 because of the better match between the predicted and real signal.

We present a number of simulations to validate MCPA for a realistic range of signal-to-noise ratios (SNR) and to show that MCPA is insensitive to local information processing. We apply MCPA to examine the inter-regional representation for natural visual stimuli in visual cortex using functional magnetic resonance imaging (fMRI) data.Specifically, we show that the interactions between regions of the visual stream (V1, V2, V3, V4, and lateral occipital cortex [LO]) are sensitive to information about individual natural images. We combine MCPA with representational similarity analysis to demonstrate that MCPA can be used to evaluate computational models and make inferences regarding the underlying neural mechanism of information transferring. To demonstrate MCPA’s applicability to electrophysiological signals and multivariate oscillatory synchrony, we use MCPA to examine the circuit-level representation for faces using intracranial electroencephalography (iEEG) data. Specifically, we show that the interaction between the occipital face area (OFA) and the fusiform face area (FFA) represents information about individual faces. These results demonstrate that MCPA can be used to probe the nature of representational space resulting from processing distributed across neural regions.

## Materials and methods

### Overview

The MCPA method consists of a learning phase and a test phase (as in machine learning, where a model is first learned, then tested). In the learning phase, the connectivity maps for each condition that characterize the pattern of shared activity between two populations is learned. In the test phase, these maps are used to generate predictions of the activity in one population based on the activity in the other population as a factor of condition and these predictions are tested against the true activity in the two populations. Similar to linear regression where one can generate a prediction for the single variable A given the single variable B based on the line that correlates A and B, MCPA employs a canonical correlation model (a generalization of multivariate linear regression) and produces a mapping model for each condition as a hyperplane that correlates multidimensional spaces A and B. Thus one can generate a prediction of the observation in multivariate space A given the observation in multivariate space B on a single trials basis. In this sense, MCPA is more analogous to a machine learning classifier combined with a multivariate extension of PPI (Friston et al., 1997) rather than being analogous to correlation-based functional connectivity measures.

The general framework of MCPA is to learn the connectivity map between the populations for each task or stimulus condition separately based on training data. Specifically, given two neural populations (referred to as A and B), the neural activity of the two populations can be represented by feature vectors in multi-dimensional spaces (Haxby et al., 2014). The actual physical meaning of the vectors would vary depending on modality, for example spike counts for a population of single unit recordings; time point features for event-related potentials (ERP) or event-related fields; time-frequency features for electroencephalography, electrocorticography (ECoG) or magnetoencephalography; or single voxel blood-oxygen-level dependent (BOLD) responses for functional magnetic resonance imaging. A mapping between A and B is calculated based on any shared information between them for each condition on the training subset of the data. This mapping can be any kind of linear transformation, such as any combination of projections, scalings, rotations, reflections, shears, or squeezes.

These mappings are then tested as to their sensitivity to the differential information being processed between cognitive conditions by determining if the neural activity can be classified based on the mappings. Specifically, for each new test data trial, the maps are used to predict the neural activity in one area based on the activity in the other area and these predictions are compared to the true condition of the data. The trained information-mapping model that fits the data better is selected and the trial is classified into the corresponding condition. This allows one to test whether the mappings were sensitive to the differential information being represented in the neural interaction in the two conditions.

The flow of the MCPA framework is demonstrated in Figure 1 and Algorithm 1.

#### Algorithm 1: Multi-Connection Pattern Analysis (MCPA)

**Input:**

training data: matrices 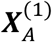 for ROI-A under condition 1, 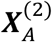 for ROI-A under condition 2, 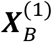 for ROI-B under condition 1, 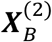 for ROI-B under condition 2 testing data: *x_A_* for observation in ROI-A, and *x_B_* for observation in ROI-B

**Output:**

Prediction of condition for observation (*x_A_*, *x_B_*).

Learning phase:

1. Apply CCA on 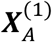 and 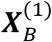 to get linear mapping function ***R***^(1)^
2. Apply CCA on 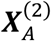 and 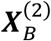 to get linear mapping function ***R***^(2)^.

Testing phase:

3 Use *x_A_* and ***R***^(1)^ to reconstruct activity in ROI-B under condition 1, which yields reconstructed data matrix 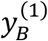.
4 Use *x_A_* and ***R***^(2)^ to reconstruct activity in ROI-B under condition 2, which yields reconstructed data matrix 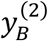.
5 Compare the correlations between the reconstructions 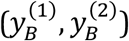 under different conditions and the real observation (*x_B_*).
6 Reverse the direction (project B to A), repeat steps 3 and 4, and compare the correlations between the reconstructions under different conditions and the real observation.
7 Assign the condition that gives maximum average correlation coefficient to the test case (*x_A_, x_B_*).

### Connectivity Map

The first phase of MCPA is to build the connectivity map between populations. The neural signal in each population can be decomposed into two parts: the part that encodes shared information, and the part that encodes non-shared local information (including any measurement noise). We assume that the parts of the neural activities that represent the shared information in the two populations are linearly correlated (though, this can easily be extended by the introduction of a non-linear kernel). The model can be described as follows

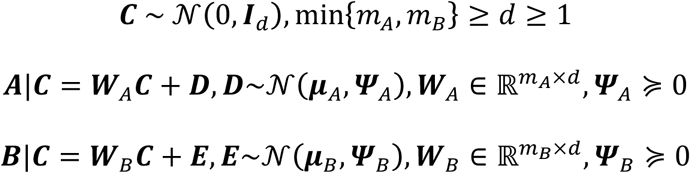
where ***C*** is the common activity, ***D*** and ***E*** are local activities, *m_A_, m_B_* are the dimensionalities of activity vector in population A and B respectively. Without loss of generality, ***μ****_A_* = ***μ****_B_* = 0 can be assumed. The activity in population A can be decomposed into shared activity ***W****_A_**C*** and local activity ***D***, while activity in B can be decomposed into shared activity ***W****_B_**C*** and local activity ***E***. The shared discriminant information only lies in the mapping matrix ***W****_A_* and ***W****_B_* since ***C*** always follows the standard multivariate normal distribution (though correlation measures that do not assume normally distributed data can also be applied with minor modifications to the calculation).

In statistics, canonical correlation analysis (CCA) is optimally designed for such a model and estimate the linear mappings (Bach and Jordan, 2005; Hardoon et al., 2004). In brief, let ***S*** be the covariance matrix

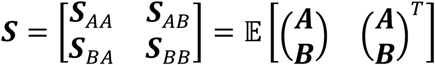

Therefore ***W****_A_* and ***W****_B_* can be estimated by solving the following eigen problem

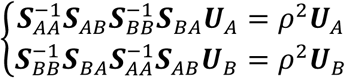
and we have

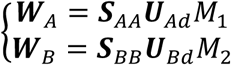
where ***U****_Ad_* and ***U****_Bd_* are the first *d* columns of canonical directions ***U****_A_* and ***U****_B_*, and ***M***_1_, 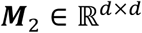 are arbitrary matrices such that 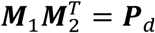, ***P****_d_* is the diagonal matrix with the first *d* elements of 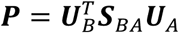. Therefore, ***M***_1_ and ***M***_2_ are just matrices used to normalize the projection of A and B onto the latent space. So ***M***_1_ and ***M***_2_ can take arbitrary value as long as 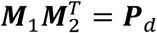, where ***P**_d_* is the diagonal matrix representing the variance along each of the *d* latent dimensions. Therefore, we can just take 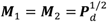.

With ***W****_A_* and ***W****_B_*, the shared information *C* can be estimated using its posterior mean 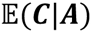 and 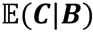, where 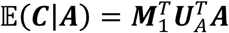 and 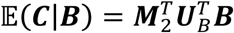. Let ***M***_1_ = ***M***_2_ and equate 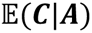 and 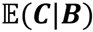, this shared information can be used as a relay to build the bidirectional mapping between A and B. Specifically, 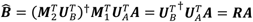 and 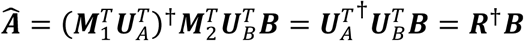, where 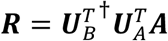.

In the first step, the connectivity map is estimated for each condition separately. If we have *n*_1_ trials in condition 1 and *n*_2_ trials in condition 2 in the training set, the training data for the two conditions are represented in matrices as 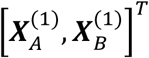 and 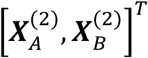 respectively, where 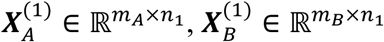 are the population activity for A and B under condition 1 respectively, and 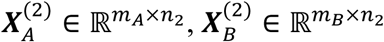 are the population activity for A and B under condition 2 respectively. The testing data vector is then represented as [***x**_A_*, ***x**_B_*]^*T*^, where 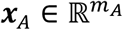 and 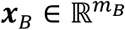 are population activities in A and B respectively. Using CCA, the estimations of the mapping matrices with respect to different conditions are ***R***^(1)^ and ***R***^(2)^.

To sum up, by building the connectivity map, a linear mapping function ***R*** is estimated from the data for each condition so that the activity of the two populations can be directly linked through bidirectional functional connectivity that captures only the shared information.

### Classification

The second phase of MCPA is a pattern classifier that takes in the activity from one population and predicts the activity in a second population based on the learned connectivity maps conditioned upon the stimulus condition or cognitive state. The testing data is classified into the condition to which the corresponding model most accurately predicts the true activity in the second population.

The activity from one population is projected to another using the learned CCA model, i.e. 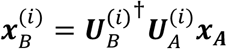. The predicted projections 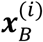 are compared to the real observation ***x_B_***, and then the testing trial is labeled to the condition where the predicted and real data match most closely. Cosine similarity (correlation) is used as the measurement of the goodness of prediction. The mapping is bidirectional, so A can be projected to B and vice versa. In practice, the similarities from the two directions are averaged in order to find the condition that gives maximum average correlation coefficient.

### Simulated experiment

To test the performance of MCPA, we used BOLD signal recorded from areas V1 and V2 to simulate shared and local activity in two populations and tested the performance of MCPA on synthetic data as a factor of the number of dimensions in each population and signal-to-noise ratio (SNR; Figure 2a). We further evaluated three control experiments to demonstrate that MCPA is insensitive to the presence or change in the local information.

For the first simulation (Figure 2a), we sampled from the empirical distribution of BOLD signal recorded from area V1 in the visual cortex and used it as the shared activity, and independently sampled signal from the empirical distributions of activity in V1 and V2 as the local unshared activity. (see fMRI method described below for experiment details). The shared activity for both conditions in population A was drawn from the empirical distribution of the first *d* principal components of V1 activity to mimic a d-dimensional normal distribution 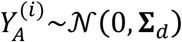 for *i* = 1,2, where **Σ**_d_ is a diagonal matrix with the jth element in the diagonal as 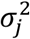. The shared activity in population B under two different conditions were generated by rotating ***Y****_A_* with different rotation matrices separately, 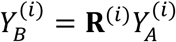, where **R**^(1)^ and **R**^(2)^ were two *d*-by-*d* random rotation matrices corresponding to the information mapping functions under condition 1 and 2 respectively, and for simplicity, **R**^(*i*)^ is orthogonal with 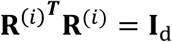. In addition to the shared activity, local activity in A and B was randomly drawn from the empirical distributions of the first *d* principal components of V1 and V2 activity respectively and multiplied by a factor of *σ* to simulate white noise 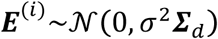.

**Figure 2.**
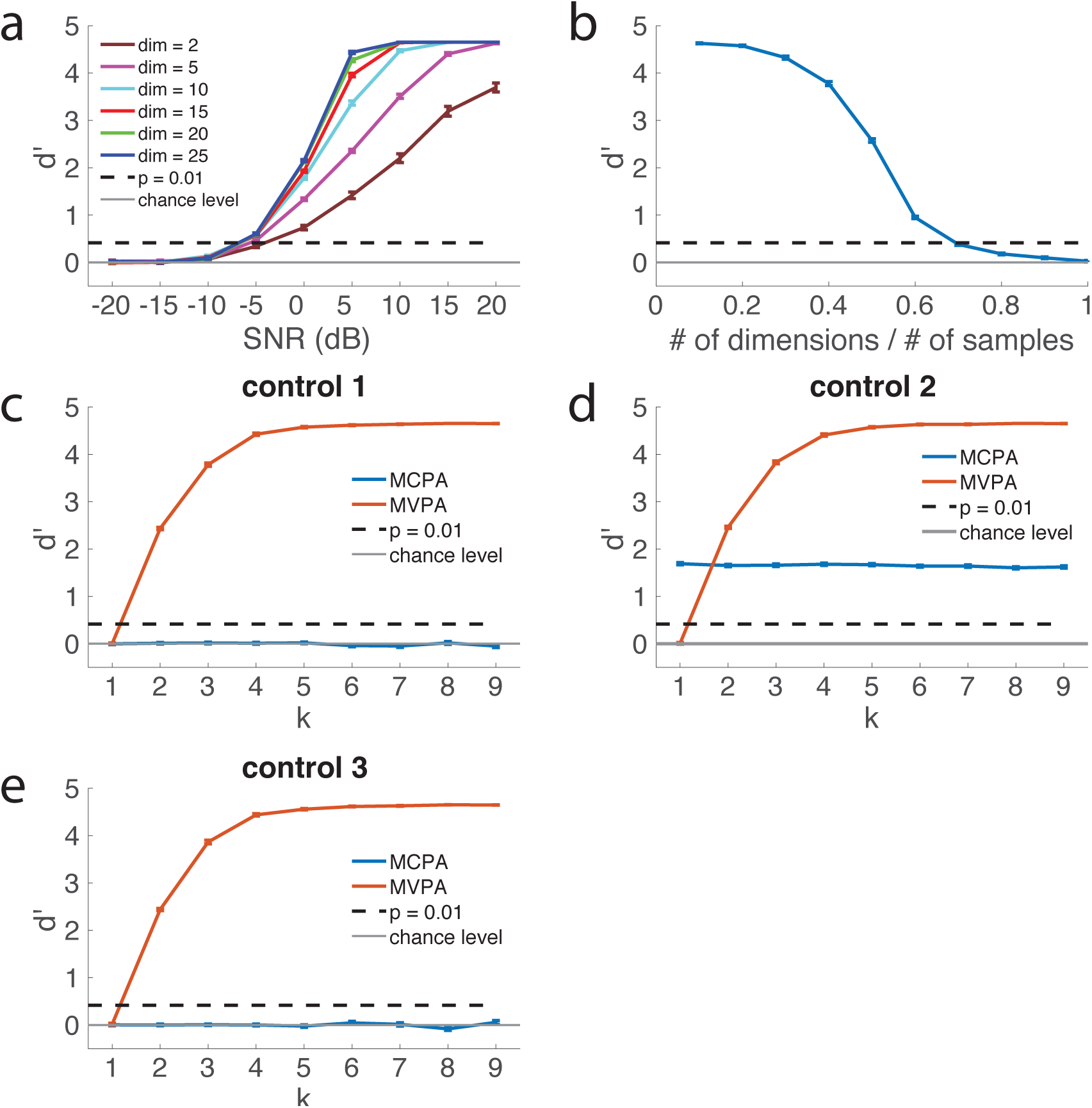
Synthetic data and control simulation experiments. The mean and standard error for 100 simulation runs are plotted. The horizontal gray line corresponds to chance level (*d*’ = 0). The dashed line (*d*’ = 0.42, corresponding accuracy 58.5%) corresponds to the chance threshold, p = 0.01, based on a permutation test. The maximum possible *d*’ = 4.65 (equivalent to 99% accuracy because the d’ for 100% accuracy is infinity). **(a)** The sensitivity of MCPA for connectivity between two populations as a factor of SNR and the number of effective dimensions in each population. MCPA was applied to synthetic data, where two conditions had different patterns of functional connectivity (measured by SNR and dimensionality). Performance of MCPA was significantly higher than chance level when SNR > -5 dB and the number of dimensions > 2. Performance of MCPA saturated to maximum when SNR > 5 dB and the number of dimensions > 10. **(b)** The robustness of MCPA to non-informative dimensions. The signal was generated in a lower dimensional manifold (# dim = 10), and *P* non-informative dimensions were added to the space. # of (training) samples per condition is fixed at 100. **(c)** The insensitivity of MCPA when there is variable local discriminant information, but no circuit-level information (control case 1). MCPA and MVPA were applied to control case 1. The SNR was fixed at 0 dB and the number of dimensions is fixed at 10 for panels b, c, and d. *k* corresponds to the ratio of the standard deviations of the two conditions in panels b, c, and d. **(d)** The insensitivity of MCPA to changes in local discriminant information with fixed circuit-level information when there is both local and circuit-level information (control case 2). **(e)** The insensitivity of MCPA to variable local discriminant information when the circuit-level activity is correlated, but does not contain circuit-level information about what is being processed (control case 3).

The two important parameters here are the dimensionality *d* and the variance *σ*^2^. SNR was used to characterize the ratio between the variance of shared activity and variance of local activity, and the logarithmic decibel scale SNR*_dB_* = −10 log_10_ (*σ*^2^) was used. To cover the wide range of possible data recorded from different brain regions and different measurement modalities, we tested the performance of MCPA with *d* ranging from 2 to 25 and SNR ranging from -20 dB to 20 dB(*σ*^2^ ranged from 0.01 to 100). Note that each of the *d* dimensions contain independent information about the conditions and have the same SNR. Thus the overall SNR does not change, but the amount of pooled information does change with *d*. For each particular setup of parameters, the rotation matrices **R**^(*i*)^ were randomly generated first, then 200 trials were randomly sampled for each condition and evenly split into training set and testing set. MCPA was trained using the training set and tested on the testing set to estimate the corresponding true positive rate (TPR) and false positive rate (FPR) for the binary classification. The sensitivity index *d*’ was then calculated as *d*′ = *Z*(*TPR*) − *Z*(*FPR*), where *Z*(*x*) is the inverse function of the cdf of standard normal distribution. This process was repeated 100 times and the mean and standard errors across these 100 simulations were calculated. Note that the only discriminant information about the two conditions is the pattern of interactions between the two populations, and neither of the two populations contains local discriminant information about the two conditions in its own activity. We further tested and confirmed this by trying to classify the local activity in populations A and B (see below). To avoid an infinity *d*’ value, with 100 testing trials, the maximum and minimum for TPR or FRP were set to be 0.99 and 0.01, which made the maximum possible *d*’ to be 4.65.

The MCPA method captures the pattern of correlation between neural activities from populations and is invariant to the discriminant information encoded in local covariance. To see this, we first take the simulation data described above and apply MVPA (naïve Bayes) to each of the two populations separately. Note that in each of the two populations, we set the two conditions to have the same mean and covariance. As a result, there should be no local discriminant information within any of the two populations alone.

#### Robustness of MCPA to non-informative dimensions

In addition to the existing simulations that evaluate the influence of SNR and informative dimensionality on the performance of MCPA, we evaluated the influence of having non-informative dimensionality on the performance of MCPA. Specifically, we simulate 10 informative dimensions and simulate *P* additional dimensions that are not informative for discrimination and apply MCPA to this simulated data without PCA. With a fixed number of 100 training samples per condition, we evaluate the performance of MCPA as a factor of the ratio between number of dimensions and the number of training samples per condition. The intuition is that, with a fixed amount of informative dimensions and fixed number of training samples, when the number of dimensions grows, the model would suffer from overfitting and the performance would decay.

#### Control simulations

For the first control simulation (Figure 2c), we fixed the dimensionality at *d* = 10 and SNR at 0 dB (*σ*^2^ = 1). For condition 1, 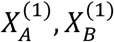 were drawn independently from the empirical distributions of the first *d* principal components of area V1 and area V2 using the corresponding empirical distributions; for condition 2, 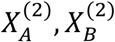 were drawn independently from the same distribution in the empirical distributions of the first *d* principal components of area V1 and area V2. Then we changed the local variance in one of the conditions. For the features in population A and B under condition 1, we used 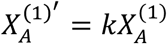 and 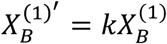 where *k* ranged from 1 to 9. Thus, in both populations, the variance of condition 1 was different from the variance of condition 2, and such difference would increase as *k* became larger. Therefore, there was no information shared between the two populations under either condition, but each of the population had discriminant information about the conditions encoded in the variance for any *k* ≠ 1.

For the second control simulation (Figure 2d), we fixed the dimensionality at 10 and SNR at 0 dB (*σ*^2^ = 1) and kept the rotation matrices of different conditions different from each other. As a result, the amount of shared discriminant information represented in the patterns of interactions stayed constant. Then we changed the local variance in one of the conditions. For the features in population A under condition 1, we used 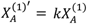, where *k* ranged from 1 to 9. Thus, population A, the variance of condition 1 was different from the variance of condition 2, and such difference would increase as *k* became larger. According to our construction of MCPA, it should only pick up the discriminant information contained in the interactions and should be insensitive to the changes in local discriminant information from any of the two populations.

For the third control simulation (Figure 2e), we introduced local discriminant information into the two populations to demonstrate that MCPA is insensitive to the presence of constantly correlated local information (Figure 2e). We fixed the dimensionality at 10 and SNR at 0 dB (*σ*^2^ = 1) and kept the rotation matrices constant for different conditions. As a result, the amount of shared discriminant information represented in the patterns of interactions was 0. Then we changed the local variance in one of the conditions. For the features in population A and B under condition 1, we used 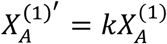 and 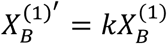, where *k* ranged from 1 to 9. Thus, in both populations, the variance of condition 1 was different from the variance of condition 2, and such difference would increase as *k* became larger. Notably, such local information was actually correlated through interactions between the populations. However, since the pattern of interaction did not vary as the condition changed, there was no discriminant information about the conditions represented in the interactions. According to our construction of MCPA, it should not pick up any discriminant information in this control case.

### Examining visual cortex coding for natural images using MCPA

#### fMRI methods

The fMRI dataset was taken from CRCNS.org (Kay et al., 2011). See (Kay et al., 2008; Naselaris et al., 2009) for details regarding subjects, stimuli, MRI parameters, data collection, and data preprocessing. In the experiment, two subjects performed passive natural image viewing tasks while BOLD signals were recorded from the brain. The experiment contains two stages: a training stage and a validation stage. In the training stage, two separate trials were recorded in each subject. In each trial, a total of 1750 images were presented to the subject, which yields a total of 3500 presentations of images (3500 = 1750 images * 2 repeats). In the validation stage, another 120 images were presented to the subject in 13 repeated trials, which yields a total of 1560 presentations (1560 = 120 images * 13 repeats). The single-trial response for each voxel was estimated using deconvolution method and used for the following analysis. The voxels were assigned to 5 visual areas (V1, V2, V3, V4, and lateral occipital [LO]) based on retinotopic mapping data from separate scans (Kay et al., 2008; Naselaris et al., 2009).

#### Categorical image classification

To control for repetition of each individual image and increase the image number being used, we used the data from the training stage for the categorical image classification. The 1750 images were manually sorted into 8 categories (animals, buildings, humans, natural scenes, textures, food, indoor scenes, and manmade objects). In order to maintain enough statistical power, only categories with more than 100 images were used in the analysis. As a result, 3 categories (food, indoor scenes, and manmade objects) were excluded.

For each pair of ROIs, namely V1-V2, V2-V3, V3-V4, and V4-LO, MCPA was applied to classify the functional connectivity patterns for each possible pair of image categories (total of 10 pairs). For each specific pair of categories, BOLD signal from all the voxels in the ROIs were used as features in MCPA. Principal Component Analysis (PCA) was used to reduce the dimensionality to *P*, where *P* corresponds to the number of PCs that capture 90% of variation in the data, which yielded between 100-200 PCs. Leave-one-trial-out cross-validation was used in order to estimate the classification accuracy. This procedure was repeated for all 10 pairs. Classification accuracy and the corresponding sensitivity index *d*’ were used to quantify the performance of MCPA.

#### Single image classification using MCPA

For single image classification the 13 repetitions of each individual image from the validation stage data was used.

For each pair of ROIs, namely V1-V2, V2-V3, V3-V4, and V4-LO, MCPA was applied to classify the functional connectivity patterns for each possible pair of images (total of 7140 pairs). For each specific pair of categories, BOLD signal from all the voxels in the ROIs were used as features in MCPA. Considering the limited number of trials in each condition, PCA was first used with the data from the training stage to reduce the representation dimensionality to 10. Because the top PCs that explain most variations may contain variance not related to the stimuli, the 10 PCs were selected from the top 50 PCs, based on maximizing the between-trial correlations for single images. As a result, we reduced the dimensionality of the validation data from more than 1000 to 10 based on the training dataset, which was completely independent from all the validation data that was used in the learning and testing stages of MCPA. Leave-one-trial-out cross-validation was then used in order to estimate the classification accuracy. This procedure was repeated for all 7140 pairs. Classification accuracy and the corresponding sensitivity index *d*’ were used to quantify the performance of MCPA.

#### MVPA analysis

MVPA was applied to classify the neural activity within each ROI (V1, V2, V3, V4, and LO) for each possible pair of categories (total of 10 pairs). The same features extracted from all the voxels within the ROI, as described above, were used in MVPA analysis. Naïve Bayes classifier was used as the linear classifier and leave-one-trial-out cross-validation was used in order to estimate the classification accuracy. This procedure was repeated for all 10 pairs. Classification accuracy and the corresponding sensitivity index *d’* were used to quantify the performance of MVPA.

#### Permutation test

Permutation testing was used to determine the significance of the classification accuracy *d’*. For each permutation, the condition labels of all the trials were randomly permuted and the same procedure as described above was used to calculate the classification accuracy (*d*’) for each permutation. The permutation was repeated for a total of 1000 times. The classification accuracy (*d*’) of each permutation was used as the test statistic and the null distribution of the test statistic was estimated using the histogram of the permutation test.

#### Representational similarity analysis

Based on the classification results, for each classification analysis, the representational dissimilarity matrix (RDM) **M** was constructed such that the *j*th element in the *i*th row, *m_ij_*, equals the dissimilarity (classification accuracy) between the condition *i* and condition *j* in the corresponding representational space defined by the analysis. Spearman’s rank correlation was used to compare representational dissimilarity matrices in order to account for outliers and non-normality in the data.

#### Psychophysiological interactions

PPI (Friston et al., 1997) was used to analyze the pattern of interactions between V1 and V4 for each pair of image categories (total of 10). The response in each ROI was extracted by taking the first principal component across all voxels. The PPI model can be written as *y* = *β*_1_*x*_1_ + *β*_2_*x*_2_ + *β*_3_*x*_3_ + *ϵ*, where *y* is the response in V4, *x*_1_is the response in V1, *x*_2_ is the categorical condition (1 or -1), and *x*_3_ is the psychophysiological interaction (*x*_3_ = *x*_1_ · *x*_2_).

#### HMAX model and connectivity patterns

The implementation of HMAX model by Serre et al. (Serre et al., 2007) was used. Each image was fed into the network and the activations in the four layers (S1, C1, S2, and C2) were recorded. At each patch size level, for image *k* (*k* = 1, 2, …, 120), the activation pattern in simple layer *i* (*i* = 1, 2) is recorded as 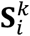, which is a square matrix with retinotopic mapping to the image space. On the other hand, the activation pattern in complex layer *i*(*i* = 1, 2) is represented as vector 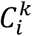 with each element representing the activation of one single unit (for C1, this is achieved by concatenating all the units in the layer into one vector). The activation of each unit in the complex layer was calculated by taking a maximum over its corresponding pool of units in the previous simple layer. For each complex unit, we recorded the location of the corresponding maximum activation simple unit. As a result, we got a *N_i_*-by-2 connectivity matrix 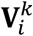 for complex layer C*i* for image *k*, where *N_i_* is the total number of units in C*i* and each row is the 2-D coordinate of the corresponding maximum activation simple unit. Thus, the connectivity pattern between simple layer S*i* and complex layer C*i* for image *k* was described by such connectivity matrix 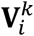 Considering all pairs of images, the RDM of the connectivity pattern **M***_i_* is calculated by taking the Frobenius norm of the difference between each pair of connectivity matrix, i.e. 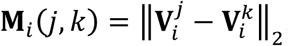.

The representation space for each single layer was then extracted by concatenating all units in the layer into one vector. The RDM of each single layer was calculated using the Euclidian distance between the corresponding activation vectors of the images.

#### Representational similarity analysis and permutation test

Permutation test was used to determine the statistical significance of the correlation between the RDM from MCPA and the RDM from HMAX. Specifically, for each pair of ROIs (i.e. V1-V2, V2-V3, V3-V4, and V4-LO), we calculated the corresponding 120-by-120 RDM for all the images from MCPA and averaged across the two subjects, noted as, **M*^ROI^*^1^** ^−^ ***^ROI^*^2^**, where *ROI*1*-ROI*2 = V1-V2, V2-V3, V3-V4, or V4-LO. Then we used the RDMs of HMAX (**M***_i_*, *i* = 1, 2) described in the previous part and calculate the Spearman’s rank correlation between **M***^ROI^***^1^** ^−^ *^ROI^***^2^** and **M***i*. As a result, we have 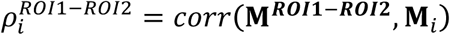. Then to compare the correlation from different layers in HMAX to MCPA, we use 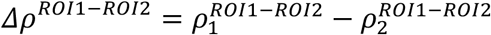 as the test statistic. For each permutation, the labels of the 120 images were randomly permuted and the above procedure was repeated. With a total of 1000 permutations, we got the empirical distribution of the test statistic for the null hypothesis that there is no difference between the two correlations. A p-value for the real test statistic can then be estimated.

### Examining OFA-FFA coding for individual faces using MCPA

#### Subject

A human subject underwent surgical placement of iEEG depth electrodes (stereotactic electroencephalography) as standard of care for surgical epilepsy localization. The subject was male, age 56. There was no evidence of epileptic activity shown on the electrodes used in this study.

The experimental protocols were approved by the Institutional Review Board of the University of Pittsburgh. Written informed consent was obtained from the participant. See supplemental methods for analysis details.

## Results

### Simulations

We used simulations to test and verify the performance and properties of MCPA on synthetic data. Specifically, synthetic data generated based on real fMRI data representing neural activity of two distinct populations and the information represented in the interaction between those populations was manipulated to construct different testing conditions.

In the first simulation, we evaluated the ability of MCPA to detect information represented in the functional connectivity pattern when it was present as a factor of the SNR and the number of dimensions of the data. The mean and standard error of the sensitivity index (*d*’) from 100 simulation runs for each particular setup (dimensionality and SNR) are shown in Figure 2a. The performance of the MCPA classifier increased when SNR or effective dimensionality increased. Classification accuracy saturated to the maximum when SNR and number of dimensions were high enough (SNR > 5 dB, dimensionality > 10). The performance of MCPA was significantly higher than chance (p < 0.01, permutation test) for SNRs above -5 dB for all cases where the dimensionality was higher than 2, when the pattern of the multivariate mapping between the activity was changed between conditions.

In addition, we examined how robust MCPA is to adding uninformative dimensions, which also changes the global SNR though non-uniformly. This simulation assesses performance of MCPA as the dimensionality of the data approaches the number of trials. In the evaluation with a fixed number of 10 informative dimension and 100 trials per condition, MCPA was shown to be highly robust to uninformative dimensions and gave significant classification accuracy until the number of dimensions approached ~70% of the number of trials (Figure 2b), at which point more than 85% of the total features are completely uninformative noise.

The first control simulation was designed to confirm that when two unconnected populations both carry local discriminant information, MCPA would not be sensitive to that piece of information. As shown in Figure 2c, MCPA did not show any significant classification accuracy above chance (*d*’ = 0) as *k* changed. On the other hand, the MVPA classifier that only took the data from local activity showed significant classification accuracy above chance level and the performance increased as local discriminant information increased.

The second control simulation was designed to test if MCPA would be insensitive to changes in local discriminant information when there was constant information coded in neural communication. Local discriminant information was injected into the populations by varying the ratio of the standard deviation (k) between the two conditions. When MVPA was applied to the local activity, increasing classification accuracy was seen as *k* became larger (Figure 2d). This result confirmed that discriminant information was indeed encoded in the local activity in the simulation. On the other hand, the performance of MCPA did not change with the level of local discriminant information (*d*’stayed around 1.65 for all cases, corresponding to accuracy = 79%), demonstrating that MCPA is only sensitive to changes in information contained in neural interactions.

The final control simulation tested whether MCPA is simply sensitive to the presence of functional connectivity between two populations *per se* or is only sensitive to whether the functional connectivity contains discriminant information. Specifically, are local discriminant information in two populations, and a correlation between their activity, sufficient for MCPA decoding? It should not be, considering that MCPA requires that the pattern of the mapping between the populations to change as a factor of the information being processed (see Figure 1). The final control simulation was designed to assess whether MCPA is sensitive to the case where two populations communicate, but in a way that would not imply distributed computational processing. Specifically, neural activity in areas A and B were simulated such that local discrimination was possible in each population and the activity of the two populations was correlated, but the interaction between them was invariant to the information being processed. Figure 2e shows that in this case MCPA did not classify the activity above chance, despite significant correlation between the regions and significant local classification (MVPA). Thus, functional connectivity between the populations is a necessary, but not sufficient, condition for MCPA decoding. Therefore, MCPA is only sensitive to the case where the mapping itself changes with respect to the information being processed, which is a test of the presence of distributed neural computation.

### Single image classification of visual cortex interactions using MCPA

To assess its performance on real neural data, MCPA was applied to Blood-oxygen-level-dependent (BOLD) fMRI measurements of human occipital visual areas, in two subjects (Subject 1 and Subject 2) during passive viewing of 13 repetitions of 120 natural images (Kay et al., 2011; Kay et al., 2008; Naselaris et al., 2009). MCPA was used for single-trial classification of these images for the interactions between V1-V2, V2-V3, V3-V4, and V4-lateral occipital (LO) cortex (e.g. 4 total region pairs * 2 subjects; see Figure 5 of Naselaris et al. (Naselaris et al., 2009) for depictions of these regions in these subjects). Across the 8 pairs of regions the mean sensitivity index (*d*’) of the single trial classification was 0.405 (SD = 0.094), with all of the pairs showing significant classification at p < 0.01 corrected for multiple comparisons (permutation test). In both subjects, MCPA classification accuracy declined going up the classic visual hierarchy. For Subject 1 and 2, the classification accuracies are shown in Table 1.

**Table 1.**
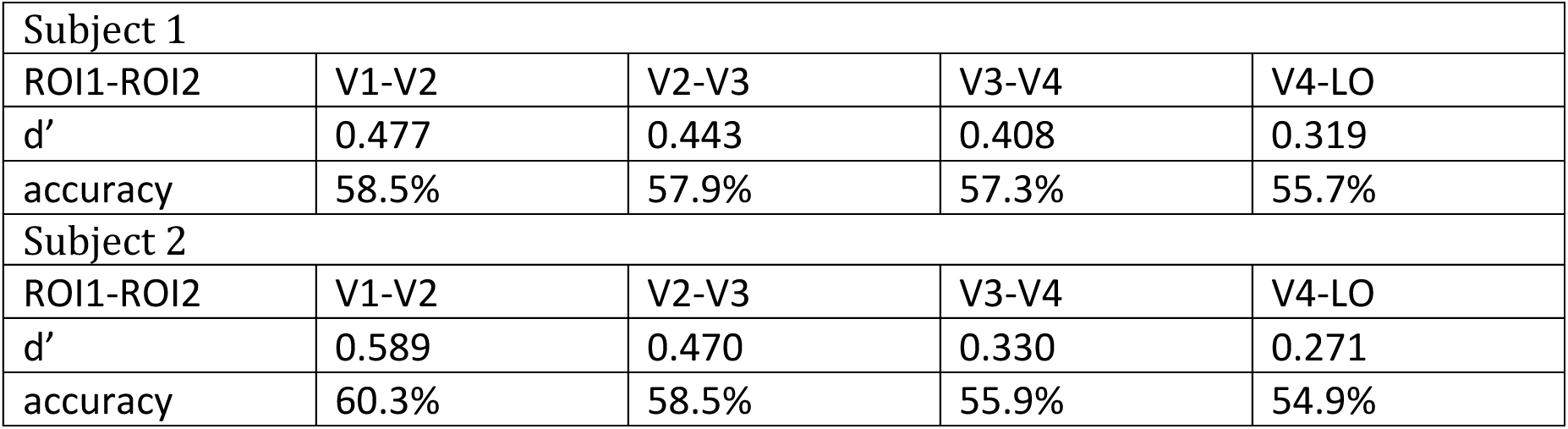
Mean d’ and classification accuracy of MCPA for Subject 1 and Subject 2 (chance level: d’ = 0, accuracy = 50%)

| Subject 1 |  |  |  |  |
| --- | --- | --- | --- | --- |
| ROI1-ROI2 | V1-V2 | V2-V3 | V3-V4 | V4-LO |
| $d'$ | 0.477 | 0.443 | 0.408 | 0.319 |
| accuracy | 58.5% | 57.9% | 57.3% | 55.7% |
| Subject 2 |  |  |  |  |
| ROI1-ROI2 | V1-V2 | V2-V3 | V3-V4 | V4-LO |
| $d'$ | 0.589 | 0.470 | 0.330 | 0.271 |
| accuracy | 60.3% | 58.5% | 55.9% | 54.9% |

### Using MCPA-based RSA to test models of between-area information transformation

One important application of MCPA is to evaluate models and test theoretical hypotheses regarding the computational operation underlying how representations are transformed from one region to another. MCPA-based representational similarity analysis (RSA) can be used to compare the representational space derived from the interaction between brain regions to representational spaces derived from the transformation of representations in computational models. To illustrate this we compare the representational space for natural images in the same fMRI dataset described above to the representational space derived from the transformation between layers of the HMAX model of the visual processing stream (Riesenhuber and Poggio, 1999; Serre et al., 2007). HMAX has four layers going from S1 to C1 to S2 to C2 along the hierarchy. The transformation of the representation between S1 and C1 (S1-C1 transformation) occurs through a local, non-linear max-pooling operation and the transformation between S2 and C2 (S2-C2 transformation) occurs through a more global non-linear max-pooling operation. We compared the representational dissimilarity matrices (RDMs) derived from these HMAX transformations to the RDMs derived from MCPA between V1-V2, V2-V3,V3-V4, and V4-LO. The transformation between C1 and S2 occurs through a passive filtering that does not give rise to an RDM because the transformation is effectively the same across all C1 representations.

As shown in Figure 3, we found that the RDM derived from the S1-C1 transformation in HMAX correlates with the V2-V3 RDM based upon MCPA of the fMRI data (mean Spearman’s rho = 0.053, p < 0.05, permutation test). Furthermore, the S1-C1 correlation to V2-V3 was significantly greater (p < 0.05, permutation test) than the S2-C2 correlation to V2-V3. The RDM derived from the S2-C2 transformation in HMAX correlates with the V4-LO RDM based upon MCPA of the fMRI data (mean Spearman’s rho = 0.112, p = 0.002, permutation test). Furthermore, the S2-C2 correlation to V4-LO was significantly greater (p < 0.01, permutation test) than the S1-C1 correlation to V4-LO. Additionally, none of the individual layers in HMAX showed a consistent significant correlation with the connectivity-based RDM from MCPA. Taken together, these results suggest that the interaction between the lower layers of the neural visual hierarchy reflects an operation more like the operation between the lower layers of the model of the visual hierarchy than between higher layers of the model. Furthermore, the interaction between higher layers of the neural visual hierarchy reflects an operation more like the operation between higher layers of the model than between lower layers of the model.

**Figure 3.**
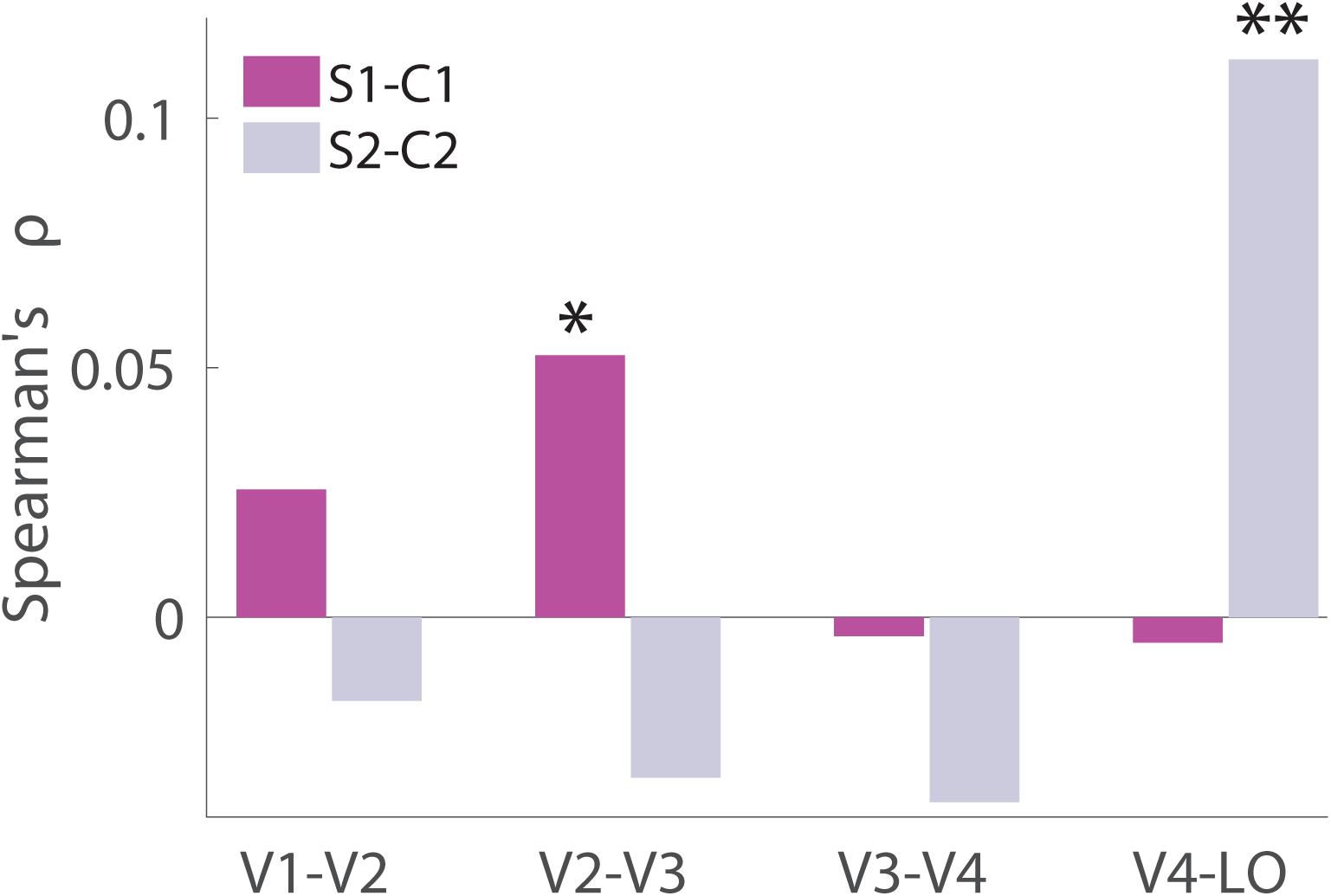
Correlating MCPA and HMAX. Correlation coefficients between the between-layer connectivity patterns in HMAX (S1-C1, and S2-C2) and the between-area connectivity patterns in fMRI data extracted by MCPA (V1-V2, V2-V3, V3-V4, and V4-LO) were plotted. The correlation was evaluated by Spearman’s rank correlation coefficients. For S1-C1, correlation peaked at V2-V3, mean Spearman’s rho = 0.053 (* p = 0.036, permutation test within each subject, and p-values were combined using Fisher’s method). For S2-C2, correlation peaked at V4-LO, mean Spearman’s rho = 0.112 (** p = 0.001, permutation test within each subject, and p-values were combined using Fisher’s method).

### Comparing the between region representation to the local representation

To assess whether the information represented in the between region interactions reflected a distinct computational process or merely reflected the representation in either of the individual areas, RSA was performed. To increase our power, we performed this RSA at the category level (animals, buildings, humans, natural scenes, and textures) based on classification accuracy rather than the single image level because the dataset contained many more repetitions per category than per image (Figure 4). This yielded a total of 16 correlations (8 MCPA-based matrices correlated with each of the two regions that contribute to each MCPA). 13 out of the 16 correlations were negative, many showing large negative correlation coefficients (see Table 2 for details, mean Spearman’s rho = -0.415, SD = 0.364). In other words, categories that were relatively easy to decode based on the activity within regions using MVPA were relatively more difficult to decode based on the shared activity between that region and the other regions in the visual stream using MCPA and vice versa (Figure 4). This negative correlation suggests that the communication between regions represents information that has not been explained aspects by local computational processes.

**Figure 4.**
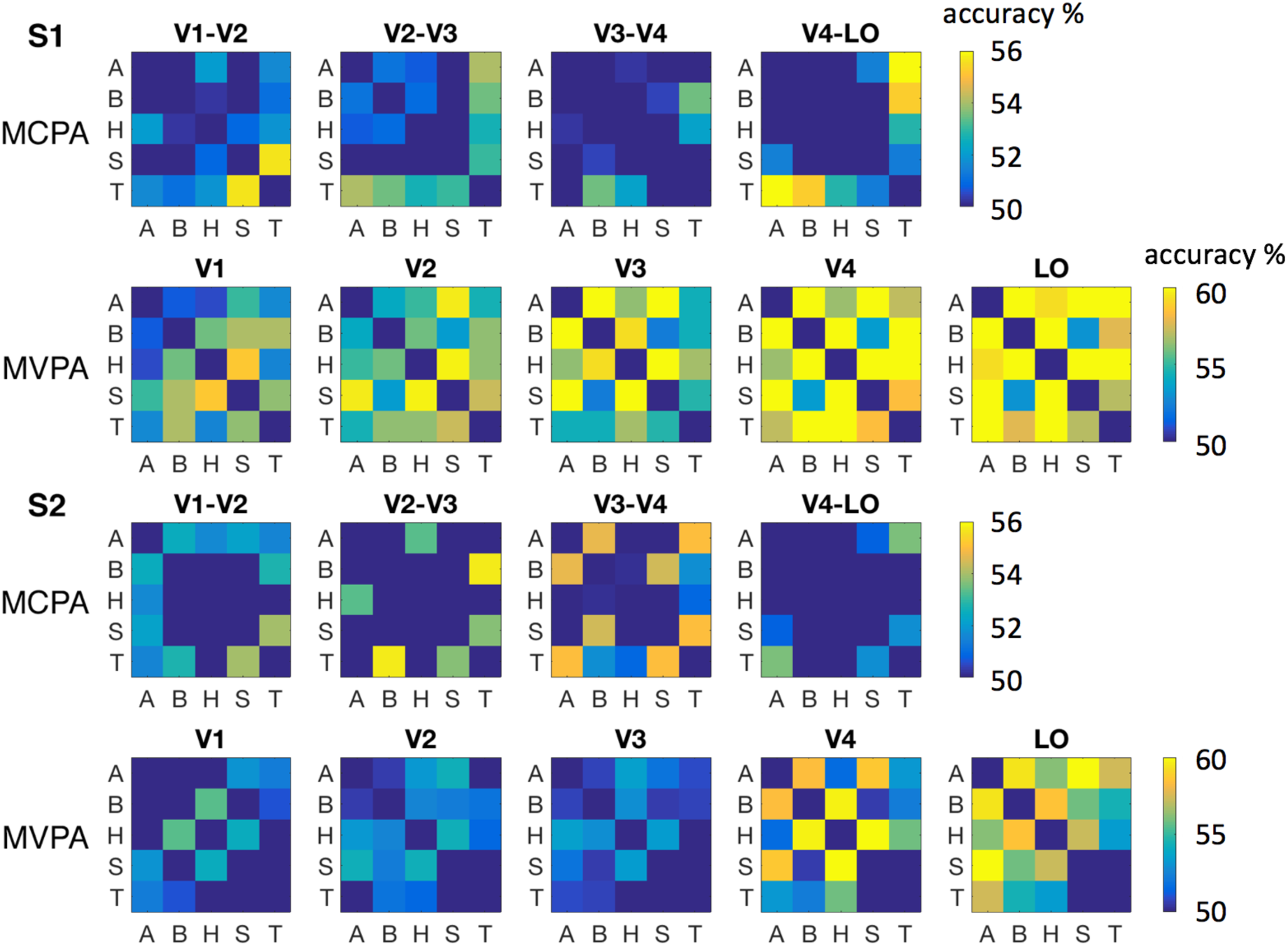
MCPA and MVPA results for fMRI categorical data. RSA results based on MCPA and MVPA for V1, V2, V3, V4, and LO from Subjects 1 and 2. Categories: A-animals, B-buildings, H-humans, S-natural scenes, T-textures. **Row 1:** RSA based on MCPA for V1-V2, V2-V3, V3-V4, and V4-LO of Subject 1, each entry represents the classification accuracy between the corresponding categories; **Row 2:** RSA based on MVPA for V1, V2, V3, V4, and LO of Subject 1, each entry represents the classification accuracy between the corresponding categories; **Row 3:** RSA based on MCPA for V1-V2, V2-V3, V3-V4, and V4-LO of Subject 2, each entry represents the classification accuracy between the corresponding categories. **Row 4:** RSA based on MVPA for V1, V2, V3, V4, and LO of Subject 2, each entry represents the classification accuracy between the corresponding categories. (chance level: accuracy = 50%).

**Table 2.**
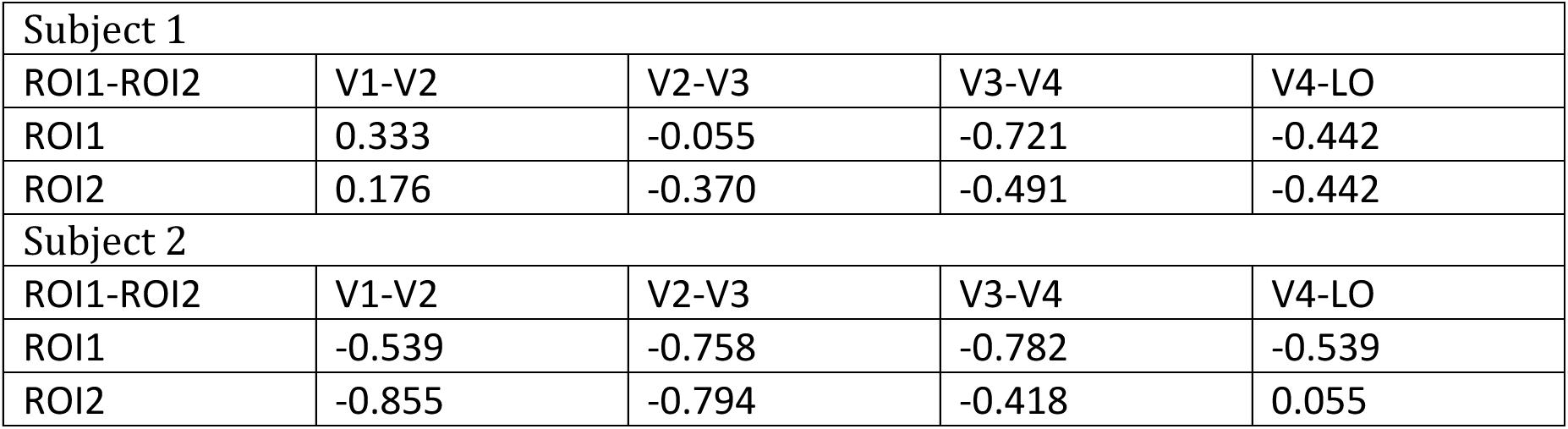
Spearman’s rank correlation coefficients between MCPA of ROI1-ROI2 and MVPA of ROI1 or ROI2 in Subjects 1 and 2.

### Comparing MCPA to PPI

To demonstrate the dominance of MCPA over classical univariate methods, we applied PPI to the same data to analyze categorical effective analysis between neighboring areas. As a comparison, 80 different pairs of categories (10 pairs of categories * 4 pairs of regions * 2 subjects) were analyzed using both PPI and MCPA. 4/80 PPI results were significant with p < 0.05 (uncorrected), while 13/80 MCPA results were significant with p < 0.05 (uncorrected). As a result, the number of significant MCPA results is significantly larger than the number of significant PPI results (p < 0.01, permutation test). Note that it is not clear how many of these 80 different pairs of categories are expected to be classifiable given that the regions examined are not category sensitive, other than LO. Thus, it is not clear if 13/80 is close to the number of category pairs that would be classifiable with perfect data or if this is a low percentage of that number, but the key point in the context of validating MCPA is that MCPA is more sensitive than univariate (PPI) methods.

### Single face identity classification of OFA-FFA interactions using MCPA

To further assess its performance on electrophysiological data, MCPA was applied on intracranial electroencephalography (iEEG) data recorded from OFA and FFA in one human epileptic patient during a visual perception task (see Figure 5a for the electrode locations). MCPA was applied in the classification between each possible pair of faces. Previous studies on the timecourse of face individuation (Ghuman et al., 2014) have demonstrated that the 250-450 ms time window is critical for the processing of face individuation information. For MCPA, as shown in Figure 5b, with a chance level of *d*’=0 and corresponding accuracy = 50%, the classification accuracy was significantly above chance level across that time window (averaged *d*’ = 0.14, mean classification accuracy 52.7%, p < 0.01, permutation test). The CCA weights for the FFA and OFA are plotted in Figure 5c, showing that 15-30 Hz in FFA and 25-40 Hz in OFA contributed most strongly to their interaction in response to individual faces, suggesting that there may be a degree of cross-frequency coupling involved in the OFA-FFA coding for faces. Using MVPA, classification accuracy was significantly above chance level across that time window in FFA (averaged *d*′ = 0.42, mean accuracy 58%, p < 0.01, permutation test), replicating previous reports (Ghuman et al., 2014), classification accuracy was also above chance level across that time window in OFA (averaged *d*’ = 0.13, mean accuracy 52.6%, p < 0.05, permutation test). In the early time window (50 – 250 ms), MCPA did not show significant classification accuracy (averaged *d*’ = 0.116, mean accuracy 51.6%, p > 0.1, permutation test).

**Figure 5.**
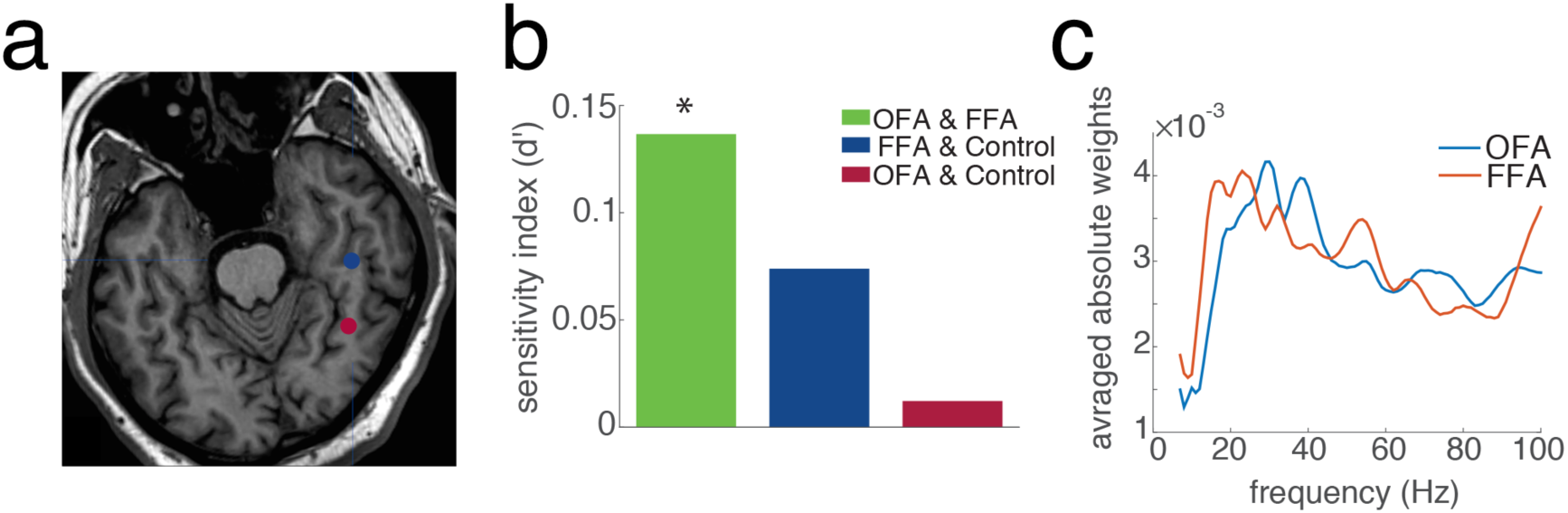
iEEG experiments and MCPA results. **(a)** Location of the electrodes of interest. The blue dot corresponds to the location of the FFA contact while the red dot corresponds to the location of the OFA contacts. **(b)** MCPA applied between (1) the OFA and FFA channels, (2) the FFA channel and the control channel, (3) the OFA channel and the control channel. The mean d’ of pairwise face classification over all 2415 pair of faces across the 200-500 ms timewindow after stimulus onset is plotted. * p < 0.01, permutation test. **(c)** Averaged absolute loading weights in the functional connectivity model of MCPA for OFA and FFA across the frequency spectrum during the time window of 250-450 ms after stimulus onset. (chance level: d’ = 0, accuracy = 50%)

As a control analysis, we took a contact outside of the fusiform gyrus that did not show face sensitivity and performed the same analysis between the control contact and the OFA and FFA contacts. As shown in Figure 5b, the averaged *d’* of MCPA between the control contact and both the OFA and FFA contacts was not significant above chance level (*d*’ = 0.074 for control & FFA, accuracy = 51.2%, d’ = 0.012 for control & OFA, accuracy = 50.3%, both p > 0.1, permutation test).

With the caveat that the effect size is small, the results support the hypothesis individual level face information is represented in the OFA-FFA interaction pattern.

## Discussion

This paper presents a novel method to assess the information represented in the patterns of interactions between two neural populations. MCPA works by learning the mapping between the activity patterns from the populations from a training data set, and then classifying the neural communication pattern using these maps in a test data set. Simulated data demonstrated that MCPA was sensitive to information represented in neural interaction for realistic SNR ranges. Furthermore, MCPA is only sensitive to the discriminant information represented through different patterns of interactions irrespective of the information encoded in the local populations. Applying this method to fMRI data demonstrated that the multivariate connectivity patterns between areas along the visual stream represent information about individual natural images. MCPA-based RSA showed that, at the category level, the representational structure of the interaction between regions is negatively correlated to the structure within each region. Furthermore, MCPA was used to test hypotheses from the HMAX model regarding the computational operation that transforms the representation between regions along the visual processing pathway. Finally, as an example with electrophysiological data, applying MCPA to iEEG data showed that the multivariate connectivity pattern between OFA and FFA represents information at the level of individual faces.

### MCPA as assessing adaptive processing

Significant discrimination within each population and significant functional connectivity between them is not sufficient to produce MCPA and indeed local classification within each population is not even necessary (Figures 2a and 2e respectively). MCPA requires the pattern of connectivity (linear correlations) between the two populations to vary across the different conditions. In other words, MCPA is sensitive to both the degree of functional connectivity in the conditions and how distinct the mappings are across conditions. As an example, if the two populations interact, but the interaction behaves like a passive linear filter, mapping the activity between the populations in a similar way in all conditions, MCPA would not be sensitive to the interaction because the mapping does not change (Figure 2e). Instead, MCPA is more akin to testing for non-constant filtering or distributed, interactive computation that behaves as a non-linear process where the nature of the interaction adapts as a factor of the information that is being processed. Recent studies demonstrate that neural populations in perceptual areas alter their response properties based on context, task demands, etc. (Gilbert and Li, 2013). These modulations of response properties suggest that lateral and long-distance interactions are adaptive and dynamic processes responsive to the type of information being processed. MCPA provides a platform for examining the role of interregional connectivity patterns in this adaptive process. Indeed, MCPA can be interpreted as testing whether distributed computational “work” is being done in the interaction between the two populations (Friston et al., 1997) and the interaction does not just reflect a passive relay of information between two encapsulated modules (Fodor, 1983).

Passive linear filters do not allow for information to be added to the representation through computational work being done in the interaction between regions. Sensitivity to this type of computation is a central appeal of fully non-linear models of neural representation and neural interactions, such as deep neural network approaches. However, these approaches often require tens of thousands or even millions of trials before they achieve good performance (Goodfellow et al., 2017), which is impractical for most neuroscientific applications. MCPA is not sensitive to multivariate non-linear interactions within conditions, but is sensitive to multivariate non-linear relationships between the interregional interaction pattern and the conditions. This is effectively a piecewise linear approximation of the underlying nonlinear function relating the condition space to the interaction pattern between regions. This restriction relative to deep neural network and other non-linear function approximation approaches allows MCPA to perform well with reasonable numbers of trials (10s of trials in our examples), which is critical for being practically useful in neuroscience. Thus, one strength of MCPA is the ability to capture some key aspects of non-linear neural computations without requiring an impractical amount of data.

### MCPA and representation space

In addition to allowing one to infer whether distributed computational work is being done in service of information processing, MCPA provides a platform for assessing its representational structure (Haxby et al., 2014). Much as MVPA has been used in representational similarity analyses to measure the structure of the representational space at the level of local neural populations (Edelman et al., 1998; Kriegeskorte, 2011; Kriegeskorte and Kievit, 2013), MCPA can be used to measure the structure of the representational space at the level of network interactions. Specifically, the representational geometry of the interaction can be mapped in terms of the similarity among the multivariate functional connectivity patterns corresponding to the brain states associated with varying input information. The representational structure can be compared to behavioral measures of the structure to make brain-behavior inferences and assess what aspects of behavior a neural interaction contributes to. It can also be compared to models of the structure to test theoretical hypotheses regarding the computational role of the neural interaction (Kriegeskorte, 2011; Kriegeskorte et al., 2008). By comparing the representational space in models to the neural representation, one can assess how well these models approximate the neural representation in both absolute and relative terms. Much the way MVPA-based RSA analyses have been used to examine these models at the level of individual brain regions (Kriegeskorte et al., 2008), RSA analyses can be used to assess how well the representation inferred by these models’ transfer functions fit the representation measured in the brain using MCPA.

The MCPA-based RSA analysis presented here relating the representational space derived from the interaction between regions of the visual processing stream to the transformation operations in HMAX is a concrete example of how MCPA can be used to test models of how representations are transformed between regions. This example also helps illustrate the underlying hypothesis being tested by MCPA: that there is a non-constant linear function that relates how the transformation of the activity between regions changes with respect to the experimental condition. A non-constant linear function is analogous to a local linear approximation of a non-linear function, as we have seen in the example of HMAX. The existence of this non-constant linear function is what allows for information to be added to the representation through distributed computational work. By comparing the MCPA-based representational space to models of this function, we can gain insight into what this transformation function might be. For example, in the case of the S1-C1 transformation HMAX, this function is a local, non-linear max-pooling operation and in the case of the S2-C2 operation it is a more global, non-linear max-pooling operation (Riesenhuber and Poggio, 1999). Furthermore, this is why MCPA could not be compared to the transformation between the C1 and S2 layers of the HMAX model because the transformation between those layers is a passive filter operation, e.g. a trivial, constant linear function relating the between layer transformation to the stimulus condition. This example suggests one mechanism by which a network with fixed structural connectivity can give rise to adaptive communication, namely through a non-linear transformation operation that are adaptive in a linear sense. In addition to testing specific hypothesis-driven transformation operations, such as the ones in HMAX, more data-driven models of the transformation operations, such as ones in deep neural network models (Yamins et al., 2014), could also be tested using the MCPA-based RSA approach.

### Relationship between MCPA and other functional connectivity/multivariate methods

These two properties of MCPA, 1) being able to assess distributed computational processing rather than just whether or not areas are communicating and 2) being able to determine the representational structure of the information being processed, set MCPA apart from previously proposed functional connectivity methods. In these previous methods the functional connectivity calculation is performed separately from the classification calculation. Specifically, either functional connectivity is first calculated using standard methods, then a model is built on the population of connectivity values and this model is tested using classification approaches (Finn et al., 2015; Richiardi et al., 2011; Rosenberg et al., 2016; Shirer et al., 2012; Wang et al., 2015) or the model is first built on the activity in each region and tested using classification approaches and the classification performance is correlated (Coutanche and Thompson-Schill, 2013; Kriegeskorte and Kievit, 2013). These methods are very useful for assessing how differences in large-scale patterns of connectivity relate to individual subject characteristics (e.g. connectome fingerprinting) in the first case and comparing the representational structure between regions in the second case. In contrast, in MCPA the model is the connectivity map and classification is done to directly test the information contained in these maps. The separation of the connectivity and classification calculations in other approaches precludes being able to assess distributed computational processes because these methods are sensitive to passive information exchange between encapsulated modules, as described above, and thus conflate passive and adaptive communication. Critically, they do not specifically probe how connectivity patterns change as a factor of condition or state, as is required to efficiently perform the representational similarity analysis in a practical manner and decode how the information processed in the interaction is encoded and organized. As a concrete example, these previous methods would not be able to compare the representational structure of the neural interaction between regions to the structure from a computational model, as was done here with fMRI.

MCPA can be roughly considered a multivariate extension of PPI with the addition of a prediction and classification framework. Compared to PPI, which is univariate, MCPA allows one to exploit the multivariate space of interaction patterns. As a result, MCPA is sensitive to aspects of information coded in interregional interactions that PPI may not be able to detect (Norman et al., 2006), for example in event-related fMRI designs where PPI is known to lack statistical power (O’Reilly et al., 2012). Indeed, in the fMRI data presented here, PPI was no better than chance in detecting interregional interactions at the visual category level, whereas MCPA was significantly better than chance. Much the way MVPA allows one to go beyond ANOVAs/t-tests in a single area/population (e.g. single trial classification, RSA, complex model testing), MCPA allows one to go beyond PPI and do these types of analyses at the level of the shared activity between regions.

The specific instantiation of MCPA presented here treats connectivity as a bi-directional linear mapping between two populations. However, the MCPA framework could be easily generalized into more complicated cases. For example, instead of using correlation-based methods like CCA, other directed functional connectivity algorithms, such as Granger causality based on an autoregressive framework, potentially using partial CCA for the time-lagged autoregressive step, could be used to examine directional interactions. This would allow one to examine time-lagged multivariate connectivity patterns to infer directionality. Additionally, kernel methods, such as kernel CCA(Hardoon et al., 2004), or deep learning methods, such as deep CCA(Andrew et al., 2013), could be applied to account for non-linear interactions. Another possible and more general framework would be to use non-parametric functional regression method to build a functional mapping between the two multidimensional spaces in the two populations. MCPA can also be expanded to look at network-level representation by implementing the multiset canonical correlation analysis, wherein the cross-correlation among multiple sets of activity patterns from different brain areas is calculated (Kettenri.Jr, 1971). MCPA could be used with a dual searchlight approach to examine whole brain communication (Kriegeskorte et al., 2006). Also, MCPA could be adapted by optimizing the CCA to find the connectivity maps that uniquely describe, or at least best separate, the conditions of interest. Furthermore, both with and without these modification, the framework of MCPA may have a number of applications outside of assessing the representational content of functional interactions in the brain, such as detecting the presence of distributed processing on a computer network, or examining genetic or proteomic interactions. MCPA is used here with fMRI BOLD signals and iEEG signal, but it can be applied to nearly any neural recording modality, including scalp electroencephalography, magnetoencephalography, multiunit firing patterns, single unit firing patterns, spike-field coherence patterns, to assess the information processed by cross-frequency coupling, etc.

### Implication from MCPA results

One caveat with the MCPA results with real data presented here is that many of the effect sizes are small. One likely reason for this is that for the decoding of individual images in fMRI and faces in iEEG the number of trials per image was very small (13 for individual images in fMRI and 15 for individual faces in iEEG). Despite the small number of trial, the classification accuracy is roughly on a par with previous exemplar-level individuation classification results using fMRI and iEEG (Ghuman et al., 2014; Nestor et al., 2011; Said et al., 2010; Skerry and Saxe, 2014). Furthermore, the HMAX-MCPA correlation is roughly on par with previously reported correlations between HMAX and single unit activity from non-human primates (Khaligh-Razavi and Kriegeskorte, 2014; Yamins et al., 2013). Given a larger number of trials, MCPA classification performance should improve. The classification performance seen here can be considered a “worst case scenario” to some extent given the low number of trials and yet performance still was not far below what has been previously reported using multivariate classification on these types of data. Nonetheless, the low effect size and small number of subjects reported here is a strong caveat to the potential neuroscientific interpretation of the fMRI and iEEG data.

The MCPA results from visual cortex show that the representational space derived from MCPA was negatively correlated to the representational space derived from MVPA from either of the local populations. This inverse relationship is consistent with the idea that the communication between regions represents information that has not been explained by local computational processes. This is supportive of models that propose coding for error propagation across the visual processing network (Rumelhart et al., 1986) or Bayesian models that suggest that visual processing occurs through iterative prediction-verification processing (Lee and Mumford, 2003). Indeed, some implementations of this class of models, interactions between regions are thought to code for prediction errors (Friston, 2010), which would predict the negative correlation seen here. More generally, these results suggest another mechanism through which a network with fixed structural connectivity can give rise to adaptive communication, namely through local or interregional recurrent interactions. With the strong caveat that these results require replication in more subjects and assessment with paradigms designed to directly test these hypotheses, this negative correlation is consistent with the hypothesis that neural interactions code for information not resolved in local computational processes.

Additional RSA analysis suggests that the transformation between lower layers of HMAX correlates with the transformation between lower layers of the ventral visual stream and the transformation between higher layers of HMAX correlates with the actual transformation between higher layers of the ventral visual stream. One question is how the representation between regions of the visual processing stream can correspond to both prediction error and the HMAX max pooling operation, as found in the two RSA analyses. One possibility is that these two operations occur at different times during visual processing, which are mixed together due to the low temporal resolution of fMRI. Indeed, HMAX is designed to model the initial feedforward sweep of visual information and the error coding is thought to occur through later recurrent and feedback processing.

The current prevalent view is that face perception is mediated by a distributed network with multiple brain areas including the OFA and FFA. Structural and functional connectivity analysis for the core network has shown that FFA is strongly connected to OFA (Gschwind et al., 2012; Ishai, 2008; Pyles et al., 2013). While these results suggest the hypothesis that face individuation may involve the interaction between these populations (and likely other face processing regions), direct evidence for this hypothesis has been lacking. Our results here support the hypothesis that individual-level facial information is not only encoded by the activity within certain brain populations, but also represented through recurrent interactions between multiple populations at a network level. This interaction was biased towards frequencies in the Beta and low Gamma bands and exhibited a degree of cross-frequency coupling. This analysis indicates that assessing cross-frequency interactions between regions is another potential application of MCPA. In addition, MCPA showed significant face individuation in approximately the 200 – 500 ms time window after stimulus onset, but did not show any significant face individuation in the early time window (50 – 200 ms after stimulus onset), which is consistent with a previous MVPA study based on iEEG recording from FFA only (Ghuman et al., 2014). This suggests that the face individuation process involves temporally synchronized, recurrent interactions between OFA and FFA and likely other nodes in the face-processing network. More broadly, the fMRI and intracranial EEG MCPA results suggest that the computational work done in service of visual processing occurs not only on the local level, but also at the level of distributed brain circuits.

## Conclusion

Previously, multivariate pattern analysis methods have been used to analyze the sensitivity to information within a certain area and functional connectivity methods have been used to assess whether or not brain networks participate in a particular process.With MCPA, the two perspectives are merged into one algorithm, which extends multivariate pattern analysis to enable the detailed examination of information sensitivity at the network level. Thus, the introduction of MCPA provides a platform for examining how computation is carried out through the interactions between different brain areas, allowing us to directly test hypotheses regarding circuit-level information processing.

## Acknowledgements

We would like to thank Kendrick Kay for sharing the fMRI dataset. We thank the patient for participating in the iEEG experiments, and Michael Ward and the epilepsy monitoring unit staff and administration for their assistance and cooperation with our research. We thank Marc Coutanche and Julie Fiez for their insightful comments and feedback on this work. This work was supported by the National Institute on Drug Abuse under award NIH R90DA023420 (to YL) and the National Institute of Mental Health under award NIH R01MH107797 and NIH R21MH103592 (to ASG). The content is solely the responsibility of the authors and does not necessarily represent the official views of the National Institutes of Health.

## Supplemental Materials

### IEEG data

#### Stimuli

In the localizer experiment, 180 images of faces (50% male), bodies (50% male), words, hammers, houses, and phase scrambled faces were used as a functional localizer. Each category contained 30 images. Phase scrambled faces were created in Matlab by taking the 2-dimensional spatial Fourier spectrum of each of the face images, extracting the phase, adding random phases, recombining the phase and amplitude, and taking the inverse 2-dimensional spatial Fourier spectrum. Each image was presented in pseudorandom order and repeated once in each session.

Faces in the individuation experiment were taken from the Karolinska Directed Emotional Faces stimulus set (Lundqvist, 1998). Frontal views and 5 different facial expressions (happy, sad, angry, fearful, and neutral) from all 70 faces (50% male) in the database were used, which yielded a total of 350 face images, each presented once in random order during a session. The patient participated in a total of 3 sessions.

All stimuli were presented on an LCD computer screen placed approximately 2 meters from participants’ heads.

#### Experimental paradigms

In the localizer experiment, each image was presented for 900 ms with 900 ms inter-trial interval during which a fixation cross was presented at the center of the screen (~ 10° x 10°of visual angle). At random, 25% of the time an image would be repeated. Participants were instructed to press a button on a button box when an image was repeated (1-back). Only the first presentations of repeated images were used in the analysis.

In the individuation experiment, each face was presented for 1500 ms with 500 ms inter-trial interval during which a fixation cross was presented at the center of the screen. Faces subtended approximately 5 degrees of visual angle in width. Subjects were instructed to report whether the face was male or female via button press on a button box.

Paradigms were programmed in Matlab™ using Psychtoolbox and custom written code.

#### Data preprocessing

The electrophysiological activity in OFA and FFA were recorded simultaneously using iEEG electrodes at 1000 Hz. They were subsequently bandpass filtered offline from 1-170 Hz using a fifth order Butterworth filter to remove slow and linear drift, the 180 Hz harmonic of the line noise, and high frequency noise. The 60 Hz line noise and the 120 Hz harmonic noise were removed using DFT filter. To reduce potential artifacts in the data, trials with maximum amplitude 5 standard deviations above the mean across the rest of the trials were eliminated. In addition, trials with a change of more than 25 μV between consecutive sampling points were eliminated. These criteria resulted in the elimination of less than 1% of trials.

As the last step of the data preprocessing, we extracted wavelet features using Morlet wavelets. The number of cycles of the wavelet was set to be 7. The entire epoch length of the data was 1500ms (-500 ~ 1000 ms relative to stimulus onset). To avoid numerical issues in MATLAB, the lowest frequency was set at 7 Hz. The wavelet features were estimated using FieldTrip™ toolbox. Finally, we took all the wavelet features at 7 - 100 Hz, with 1 Hz steps, at every 10 ms as features, which yielded a 94-dimensional feature vector at every time point. All the wavelets were normalized to the baseline by subtracting the mean value and divided by the standard deviation of the data from 350ms to 50ms before stimulus onset.

#### Electrode selection

Face sensitive electrodes were selected based on anatomical and functional considerations. Electrodes of interest were restricted to those that were located in or near the fusiform gyrus or inferior occipital cortex. In addition, MVPA was used to functionally select the electrodes that showed sensitivity to faces, comparing to other conditions in the localizer experiment. Specifically, electrodes were selected such that their peak 6-way classification *d’* score (see below for how this was calculated) exceeded 1 (*p* < 0.001 based on a permutation test, as described below) and the event related potential (ERP) for faces was larger than the ERP for the other non-face object categories.

There were 12 contacts on a depth electrode on the ventral temporal lobe extending along the anterior-posterior axis. Among all the contacts, only three (the 1^st^, 6^th^ and 7^th^ contacts, see figure 3a for the location of these contacts) satisfied the criterion described above (see Figure S1 for *d*’ timecourses from all contacts on the depth electrode). The first contact was near the mid-fusiform gyrus while the other two were near posterior end of the fusiform gyrus/anterior end of the inferior occipital cortex. Hence we used the data from the first electrode as FFA signal and the averaged data across the 6^th^ and 7^th^ electrodes as the OFA signal (see Figure S2 for averaged ERP data in the two areas). The post-operative structural MRI scan did not allow us to carefully distinguish the precise localization of the “OFA” electrodes and it may be that these electrodes are in fact in the posterior fusiform and properly labeled “FFA-1” according to the recent nomenclature introduced by Weiner et al. (Weiner et al., 2010). However, considering OFA and FFA-1 are contiguous with one another and it has not been determined what, if any, functional distinction there is between the two, we use “OFA” for the label of the electrodes out of convenience.

### Examining OFA-FFA coding for individual faces using MCPA

#### MCPA Analysis

MCPA was applied to classify the OFA-FFA connectivity for each possible pair of faces (total of 2415 pairs). For each specific pair of faces, averaged wavelet features within a 50 ms time window were used as features in MCPA. Principal Component Analysis (PCA) was used to reduce the dimensionality from 94 to P, where *P* corresponds to the number of PCs that capture 95% of variation in the data, the typical value of *P* is around 7~8. Leave-one-trial-out cross-validation was used in order to estimate the classification accuracy. This procedure was repeated for all 2415 pairs and all time windows slid with 10 ms step between 0 and 600ms after stimulus onset. Similar to previous simulations, *d’* was used to quantify the performance of MCPA.

Permutation test was used to determine the significance of the *d’* timecourse of MCPA (Maris and Oostenveld, 2007). During each permutation, the condition labels of all the trials were randomly permuted and the same procedure as described above was used to calculate the timecourse of *d’* for each permutation. The permutation was repeated for a total of 1000 times. The mean *d’* during 200-500 ms of each permutation was used as the test statistic and the null distribution of the test statistic was estimated using the histogram of the permutation test. The time window 200-500 ms was chosen based on the fact that the sensitivity of facial identity was only presented in OFA and FFA roughly 200 -500 ms after stimulus onset. (Ghuman et al., 2014)

### Testing significance of CCA models

The significance of MCPA relies on two factors: the presence of functional connectivity and the discriminant information in the connectivity patterns. Both are necessary conditions for the significance of MCPA. Here we would like to evaluate the significance of the CCA models in order to further support the MCPA results.

For categorical fMRI data, we have enough repetitions to perform parametric test. We directly performe significance test with Wilk’s lambda, and the interactions between all 4 pairs of regions for all 5 categories are significant in both subjects (p < 1e-5 for all cases).

For the single image case in fMRI, since for each condition we have only 13 repetitions, it is not reliable to directly use Wilk’s lambda and the Chi-square approximation. Therefore, we use permutation test and compute the averaged canonical correlation across all conditions as the test statistic. In all pair of ROIs in both subjects, we have p < 0.01 with 1000 permutations.

Similarly, for face individuation in iEEG data, we also use permutation test and compute the averaged canonical correlation across all conditions as the test statistic. As a result, the canonical correlation between FFA and OFA electrodes are significant with p < 0.01 based on 1000 permutations.

---

**Figure S1:**
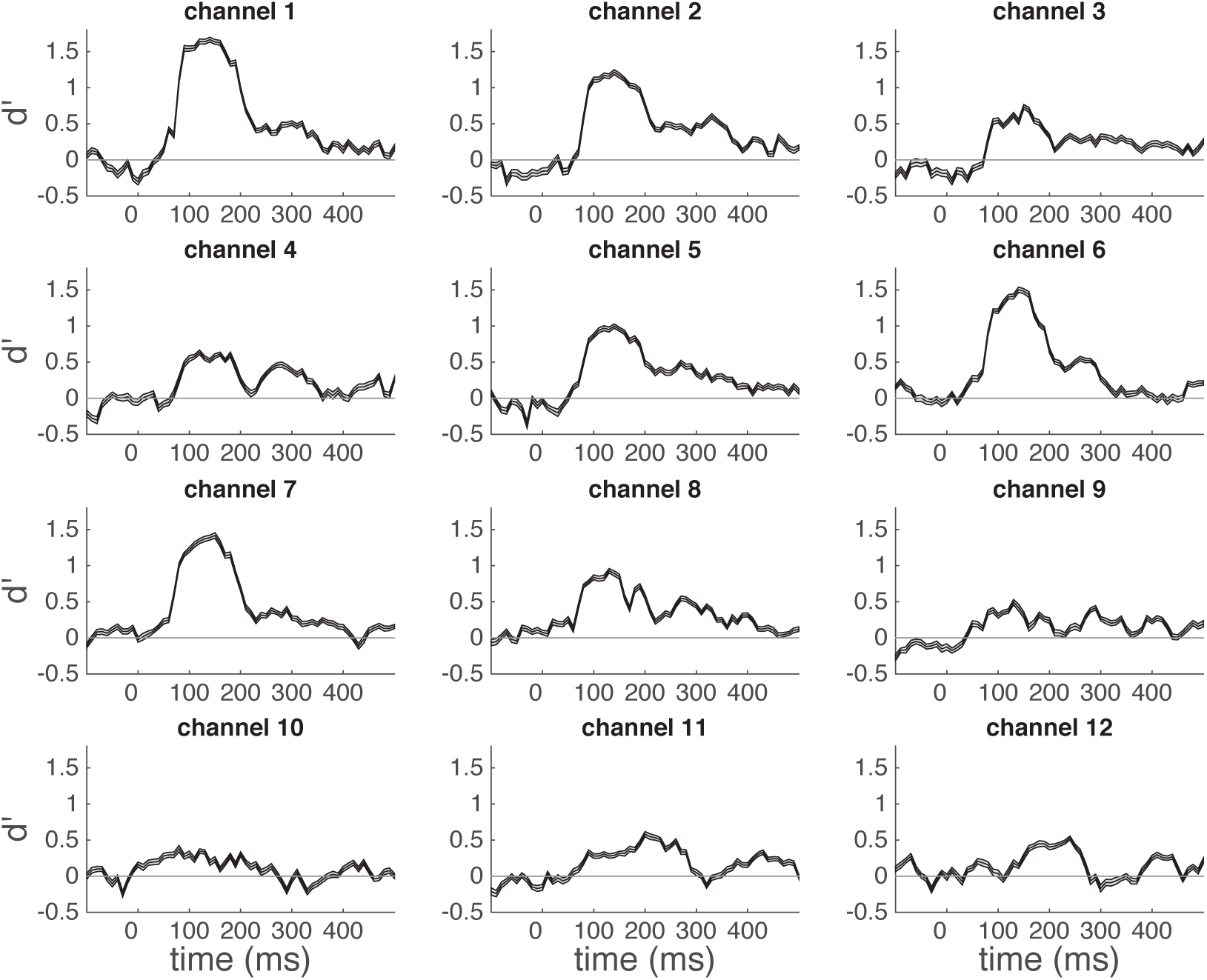
Single electrode face sensitivity. Time course of face categorical sensitivity in each single electrode measured by sensitivity index *d’* (mean *d’* plotted against the beginning of the 100 ms sliding window). The classifier uses time-windowed ERP signal from a single electrode (window length = 100 ms) as input features (See Methods for details). Horizontal grey line indicates chance level (*d’* = 0). The channels are labeled 1-12 from anterior to posterior. Electrodes were chosen based on the criteria that peak d’ be above 1 (p<0.001, channels 1, 6, and 7). Channel number 1 was used as the FFA electrode and channels 6 and 7 were used for the OFA electrodes based on their anatomical locations.

**Figure S2:**
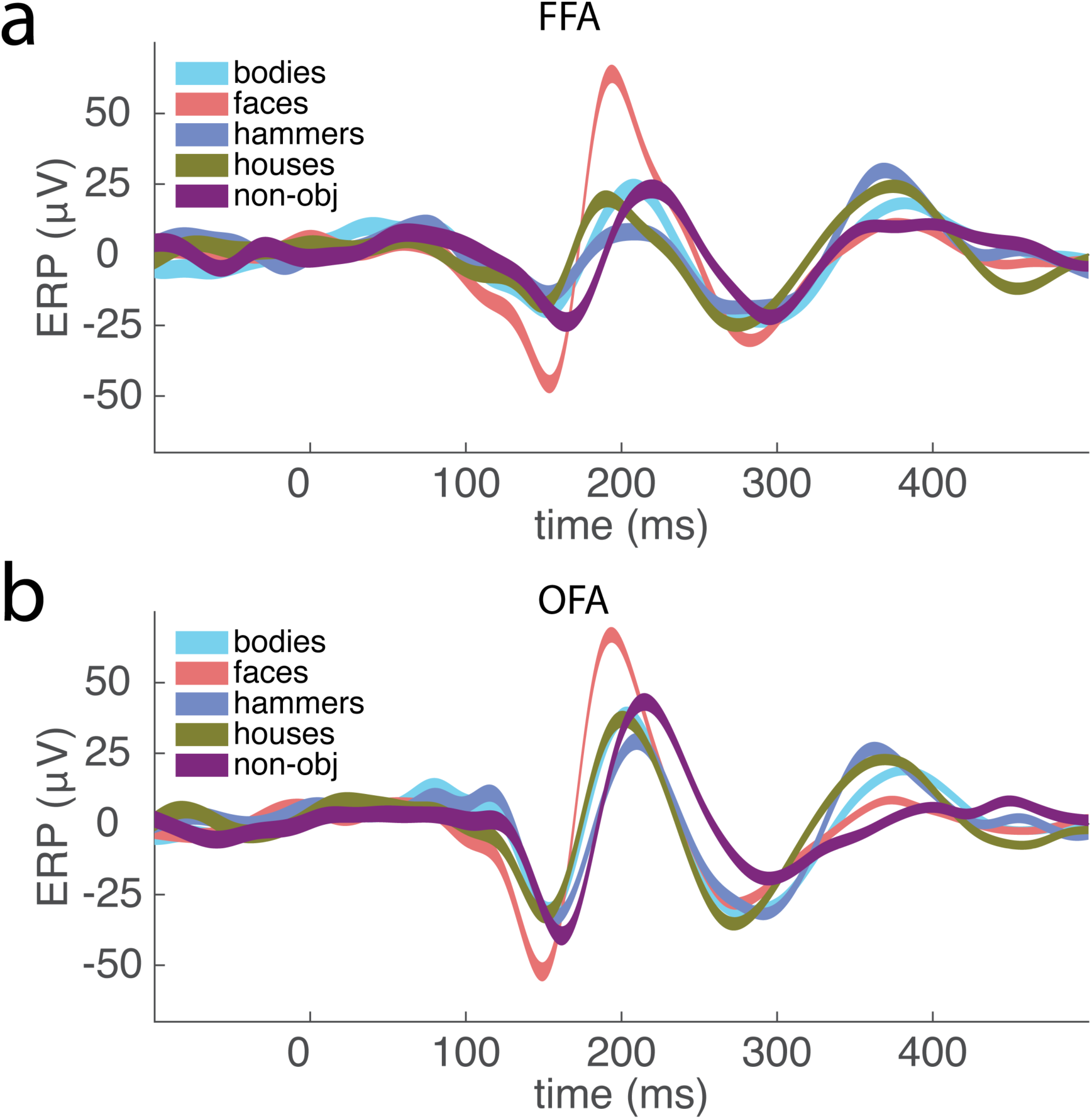
Face selectivity in FFA and OFA. Averaged ERP response recorded from FFA and OFA contacts for each category during the localizer task. The colored area corresponds to the standard error.

